# Ste5 Membrane Localization Allows MAPK Pathway Signaling *in trans* Between Kinases on Separate Scaffold Molecules

**DOI:** 10.1101/673855

**Authors:** Rachel E. Lamson, Matthew J. Winters, Peter M. Pryciak

## Abstract

The MAP kinase cascade is a ubiquitous eukaryotic signaling module that can be controlled by a diverse group of scaffold proteins. In budding yeast, activation of the mating MAP kinase cascade involves regulated membrane recruitment of the archetypal scaffold protein Ste5. This event promotes activation of the first kinase, but it also enhances subsequent signal propagation through the remainder of the cascade. By studying this latter effect, we find that membrane recruitment promotes signaling *in trans* between kinases on separate Ste5 molecules. First, *trans* signaling requires all Ste5 domains that mediate membrane recruitment, including both protein-binding and membrane-binding domains. Second, artificial membrane tethering of Ste5 can drive *trans* signaling, bypassing the need for native localization domains. Third, *trans* signaling can occur even if the first kinase does not bind the scaffold but instead is localized independently to the plasma membrane. Moreover, the *trans* signaling reaction allowed us to separate Ste5 into distinct functional domains, and then achieve normal regulation of signal output by tethering one domain to the membrane and stimulating membrane recruitment of the other. Overall, the results support a heterogeneous “ensemble” model of signaling in which scaffolds need not organize multiprotein complexes but instead can serve as binding sinks that co-concentrate enzymes and substrates at specific subcellular locales. These properties relax assembly constraints for scaffold proteins, increase regulatory flexibility, and can facilitate both natural evolution and artificial design of new signaling proteins and pathways.

## INTRODUCTION

Most signal transduction pathways begin at the plasma membrane. In addition to the unique role of the plasma membrane as the interface between the cell exterior and interior, localization of signaling proteins to the plasma membrane has the potential to influence their reactions by altering their access to activators, substrates, and cofactors. In principle, such effects can emanate from increases in protein concentration as a result of colocalization [1, 2]. These issues are relevant in the mating pathway of budding yeast, a model system for investigating eukaryotic signaling mechanisms [3, 4]. In this pathway (Figure 1A), extracellular mating pheromones are detected by a transmembrane G protein-coupled receptor (GPCR), which triggers dissociation of a heterotrimeric G protein (Gαβγ). The liberated Gβγ dimer then activates a downstream MAP kinase (MAPK) cascade in a manner that requires a crucial intermediary, Ste5, a multi-domain scaffold protein with binding domains for Gβγ, membrane phospholipids, and pathway kinases (Figure 1A-B). Signal transmission involves dramatic changes in subcellular localization: when Gβγ is activated, it triggers plasma membrane recruitment of Ste5, which thereby mediates membrane localization of its associated kinases [5-8]. The proper membrane localization of Ste5 depends on the concerted action of multiple binding motifs (Figure 1B), including a RING-H2 domain that interacts with Gβγ, a short membrane-binding peptide called the PM domain, and phospholipid-binding sequences within a larger PH domain [9-11]. Essential interactions with pathway kinases are mediated by two globular regions, the PH domain and the VWA domain (Figure 1B).

**FIGURE 1:**
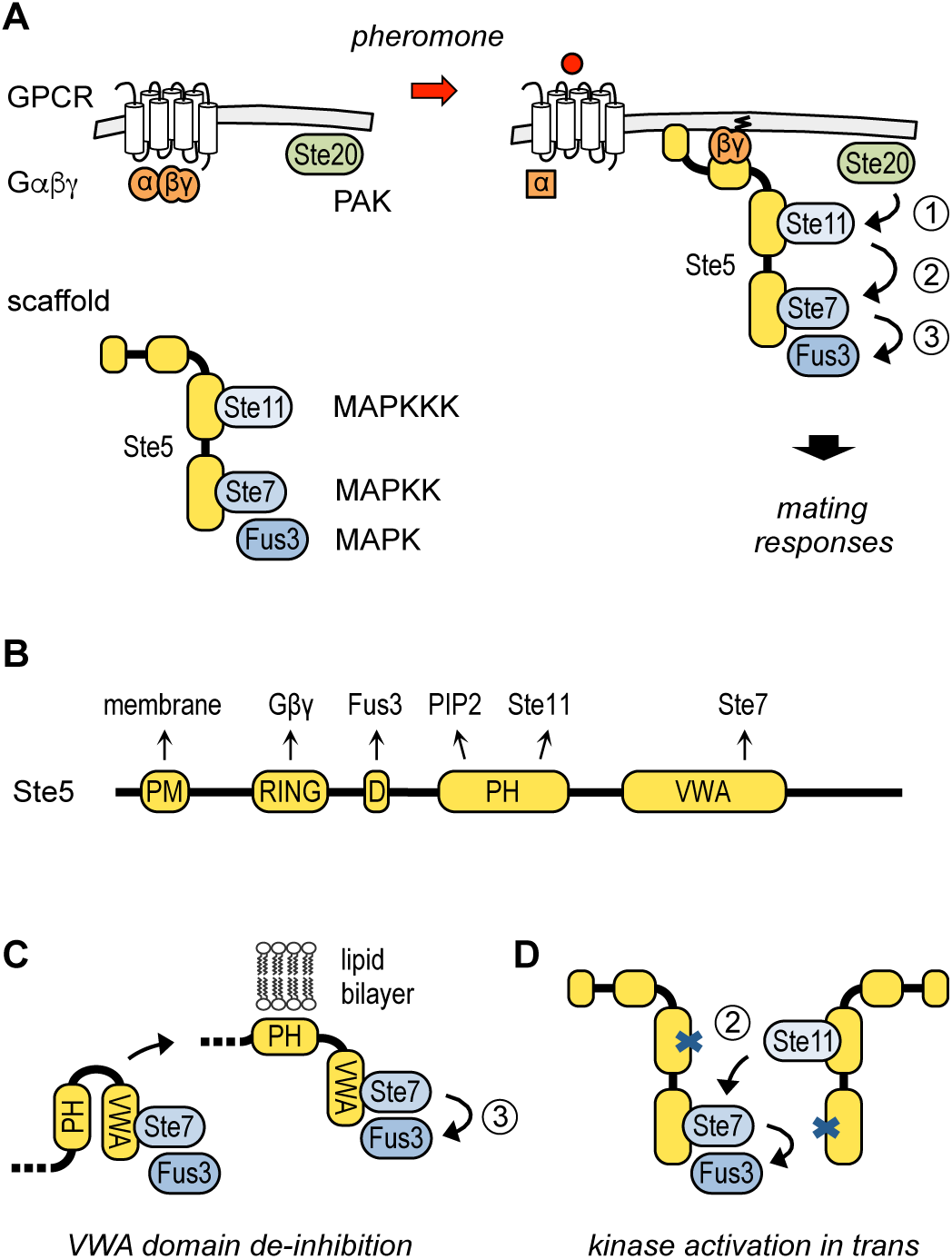
Pheromone response pathway activation and role of the scaffold protein. (A) The pheromone response pathway. Binding of pheromone to the GPCR triggers dissociation of the G protein heterotrimer (Gαβγ). The free Gβγ dimer then stimulates membrane recruitment of the scaffold protein, Ste5. A sequential cascade of three kinase activation steps ultimately activates the MAPK Fus3 (and its semi-redundant paralog, Kss1, not depicted), which induces downstream mating responses. (B) Domain structure of Ste5, with binding targets of each region indicated above [10, 11, 19, 22, 47-49. Note that the Fus3-binding motif (“D”) allows Fus3 to trigger negative feedback but is not required for positive signaling [49]; instead, Fus3 must bind directly to Ste7 [26], as implied in panel A. (C) Lipid bilayer contact releases the Ste5 VWA domain from inhibition by the PH domain [16], allowing the VWA domain to promote the final Ste7 → Fus3 step by inducing Fus3 to be a receptive Ste7 substrate [22]. (D) Kinase activation can occur *in trans*, based on complementation between two Ste5 mutants that are each defective in binding one kinase [17, 18].

The binding and membrane recruitment of Ste5 by activated Gβγ has multiple effects on signaling through the downstream pathway. First, it helps initiate signaling by allowing membrane-bound Ste20 molecules to activate the first Ste5-associated kinase, the MAP kinase kinase kinase (MAPKKK) Ste11 (Figure 1A, step 1) [5, 12]. Second, it enhances the efficiency of signal propagation from Ste11 through the remainder of the kinase cascade (Figure 1A, steps 2-3). Specifically, if Ste11 is constitutively pre-activated by mutation (thus bypassing Ste20), pathway output remains low until cells are treated with pheromone or Ste5 is artificially localized to the plasma membrane [13, 14]. An analogous effect occurs in the mammalian Raf-MEK-ERK cascade, where membrane localization can convert low Raf activity into high pathway output [15]. This ability of membrane localization to stimulate steps in the middle of the pathway, rather than just at the top, can help prevent crosstalk from other pathways that use shared components [13, 16]. It can also help shape the input-output properties of the pathway, by favoring a graded response to increasing levels of stimulus [14]. Yet the molecular mechanisms for these effects on signal propagation are only partly understood. As one contributor, membrane contact is thought to release the Ste5 VWA domain from inhibition by the PH domain (Figure 1C), and thus promote the final step in which the MAP kinase kinase (MAPKK) Ste7 activates the MAPK Fus3 [16]. The current study investigates whether membrane localization can also promote the previous step, in which Ste11 activates Ste7.

Although scaffold proteins are usually assumed to promote reactions between proteins bound to one scaffold molecule, previous observations suggest that the Ste11 → Ste7 activation step can occur with the two kinases bound to different molecules of Ste5 (Figure 1D). Namely, Ste5 mutants that cannot bind either Ste11 or Ste7, and hence are non-functional when expressed alone, will complement each other when co-expressed in the same cell [17, 18]. This finding implies that Ste11 can phosphorylate Ste7 molecules on a separate scaffold, or “*in trans*”. At the time, this phenomenon was hypothesized to reflect dimerization of Ste5 [17, 18]. Moreover, because *trans* signaling required the RING-H2 domain, it was suggested that this domain mediates dimerization. However, these studies were performed prior to any published knowledge about subcellular localization of Ste5 and the pathway kinases, or their Gβγ-regulated membrane association. In retrospect, we wondered if *trans* signaling might be a more general consequence of membrane colocalization that could be broadly applicable to a variety of signaling proteins and pathways. Therefore, we revisited this phenomenon to better understand its mechanistic implications. Here, we report that *trans* signaling requires all membrane localization sequences, and that membrane localization is indeed both necessary and sufficient for *trans* signaling. Moreover, the capacity for *trans* signaling allows the distinct kinase-binding domains of Ste5 to be separated and localized independently, and yet the pathway remains controllable by stimulus-mediated membrane recruitment of either domain. By colocalizing and concentrating reactants in a reduced subcellular volume, membrane localization can allow Ste5 to enhance signal propagation without any individual scaffold molecules being fully occupied with kinases. These properties provide functional flexibility that may foster rapid evolution of scaffold proteins, and they have broad implications for membrane-localized signaling in diverse pathways.

## RESULTS

### *Trans* signaling requires membrane localization

We used the original inter-allelic complementation assay [17, 18] to test which Ste5 domains were required for *trans* signaling. In particular, we sought to determine if *trans* signaling requires a unique dimerization motif or all sequences involved in membrane recruitment. We started with two Ste5 mutants that have defects in kinase binding (Figure 2A) [18, 19]. One mutant, Ste5-I504T, harbors a mutation in the PH domain that disrupts binding to Ste11. Another mutant, Ste5-V763A S861P (*aka* “VASP” [20, 21]), harbors a mutation in the VWA domain that disrupts binding to Ste7. These were compared to mutants with defects in membrane recruitment [5, 9-11] (Figure 2A): ΔPM (missing the N-terminal membrane-binding motif), ΔRING (missing the Gβγ-binding RING-H2 domain), Gβγ* (missing residues 152-173, which disrupts Gβγ binding but not other RING-H2 interactions), or PH* (mutations R407S K411S in the PH domain, which disrupt membrane interaction). Whereas the two mutants with kinase-binding defects could complement each other, they could not be complemented by any of the four membrane localization mutants (Figures 2B, S1A). For further tests we incorporated the localization mutations into the Ste5-VASP mutant, and measured signaling output by assaying transcription and MAPK activation. By any assay, the localization mutations disrupted the ability of Ste5-VASP to complement Ste5-I504T (Figures 2C-D, S1B). To counter the possibility that the localization sequences might have cryptic dimerization functions, we replaced the PM domain with an unrelated membrane-binding motif [10] (Figure 2A): the PH domain from PLCδ (in one or two copies). This replacement restored *trans* signaling (Figures 2C-D, S1B), arguing that the deficiency in the ΔPM mutant can be explained by its localization defect. Therefore, *trans* signaling requires all localization sequences in Ste5, and hence the role of the RING-H2 domain is not unique but instead exemplifies a more general need for membrane localization. Notably, the quantitative assays indicated that *trans* signaling is an efficient reaction (rather than a rare event), as the pathway output was within 25-50% of that obtained with intact Ste5 (Figures 2C, S1A,C).

**FIGURE 2:**
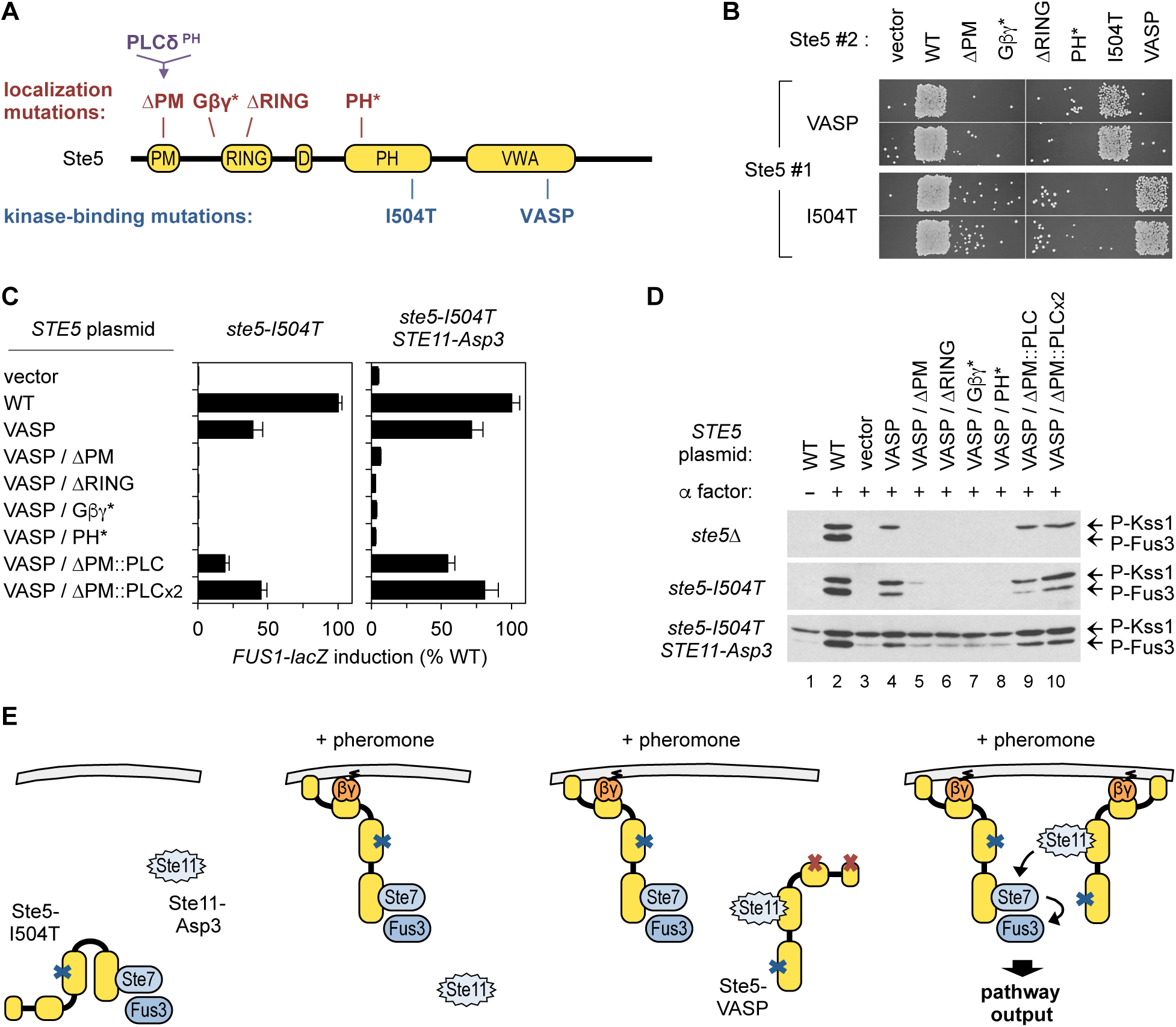
*Trans* signaling requires all membrane localization sequences in Ste5. (A) Positions of Ste5 mutations that affect either membrane localization or kinase binding. In some experiments, the Ste5 PM domain is replaced with the PH domain from mammalian PLCδ (PLCδ^PH^), which restores membrane localization [10]. (B) Patch mating tests (shown in duplicate) of complementation between mutants with defects in kinase binding versus localization. Strains (PPY1974, PPY1975) with VASP or I504T mutations at the genomic locus (Ste5 #1) harbored *STE5-myc*_*13*_ plasmids (Ste5 #2). Also see Figure S1A. (C) Transcriptional induction (*FUS1-lacZ*) in *ste5-I504T* cells (PPY1975), ± *STE11-Asp3*, harboring *STE5-HA*_*3*_ variants, treated with α factor (5 µM, 2 hr). Bars, mean ± SD (n = 3). (D) MAPK phosphorylation in strains (PPY2032, PPY1975) harboring *STE5-HA*_*3*_ variants ±*STE11-Asp3*, treated ± α factor (5 µM, 15 min). Also see Figure S1B. (E) Interpretation of findings. Ste11-Asp3 is depicted with a spiky outline to denote its pre-activated state. In Ste5-I504T cells, Ste11-Asp3 yields only low Fus3 activation and pathway output, ± pheromone stimulation (left panels). Adding Ste5-VASP allows signaling *in trans*, but only if membrane localization sequences are intact (right panels).

The MAPK phosphorylation assays provided additional insights into which signaling steps were disrupted by the localization mutations. As described below, our analyses suggested that both the Ste20 → Ste11 and Ste11 → Ste7 steps were affected. When Ste5-VASP and Ste5-I504T were coexpressed, pheromone stimulated phosphorylation of both Fus3 and its semi-redundant paralog, Kss1 (Figure 2D, middle, lane 4). But when Ste5-VASP was expressed alone, pheromone could activate only Kss1 and not Fus3 (Figure 2D, top, lane 4). As reported previously [20, 21], this behavior implies that Ste5-VASP is competent to mediate activation of Ste11 but cannot promote the subsequent reactions; consequently, the activated Ste11 can dissociate from Ste5 to weakly drive these reactions “off scaffold”, leading to weak activation of Kss1 but not Fus3 (which requires the VWA domain [22]). This Kss1 phosphorylation provides a useful proxy for the successful activation of Ste11 by Ste5-VASP. Notably, it was disrupted by all four localization mutations (Figure 2D, top and middle, lanes 5-8) suggesting that they prevent Ste11 from being activated by Ste20 at the plasma membrane. In principle, this defect in Ste11 activation could suffice to explain their failure to complement Ste5-I504T. Thus, to address whether subsequent steps were also affected, we used a pre-activated form of Ste11, called Ste11-Asp3 (which contains Asp replacements at three activating phosphorylation sites) [12]. This Ste11-Asp3 mutant bypasses its upstream activator, Ste20, yet pathway output is still regulated by pheromone [13, 14].

The experiments with Ste11-Asp3 (Figures 2C, right, and 2D, bottom) revealed several noteworthy points, which ultimately suggest that membrane colocalization can promote the Ste11 → Ste7 reaction. First, in the absence of pheromone, elevated P-Kss1 levels confirmed the constitutive activity of Ste11-Asp3, while the low levels of P-Fus3 confirmed that this response still required pheromone stimulation (Figure 2D, bottom, lanes 1-2). Second, the Ste5-I504T mutant alone could not mediate the pheromone-induced increase in transcription and P-Fus3 (Figure 2C, right, and Figure 2D, bottom, lane 3). This indicates that the stimulatory effect of pheromone cannot be explained simply by relieving inhibition of the VWA domain (in the Ste5-I504T protein), which would have been expected if Ste7 were fully activated by Ste11-Asp3 prior to pheromone addition. Third, the addition of Ste5-VASP restored activation of Fus3 and transcription, but this required that the membrane localization sequences were intact (Figure 2D, bottom, lanes 4-10). This behavior suggests a need to co-localize Ste5-I504T and Ste5-VASP molecules at the membrane, in order to increase the mutual proximity of Ste11-Asp3 and Ste7, and thereby promote the MAPKKK → MAPKK reaction *in trans* (Figure 2E). In other words, these results show that the role of membrane localization in *trans* signaling cannot be explained solely by a need to activate Ste11 (which is bypassed by Ste11-Asp3) or by a need to de-repress the VWA domain (which should occur for Ste5-I504T in the absence of Ste5-VASP), and instead they support the view that the Ste11 → Ste7 reaction is promoted by the co-recruitment of the two distinct scaffold molecules.

### Membrane localization of kinase-binding domains permits *trans* signaling

To pursue the implications of our findings further, we asked if membrane localization could be sufficient for *trans* signaling (Figure 3A). Thus, we used a form of Ste5 that is tethered to the plasma membrane by a C-terminal transmembrane domain (CTM), under control of an inducible promoter (*P*_*GAL1*_). Previous studies showed that expression of this Ste5-CTM fusion protein could activate signaling and bypass the need for pheromone, receptor, and Gβγ [5]. Not surprisingly, variants of this Ste5-CTM fusion that harbored either of the kinase-binding mutations (I504T or VASP) were defective when expressed alone (Figure 3B). However, co-expression of the two membrane-tethered mutants restored signaling (Figures 3B), and with an efficiency similar to that described earlier when *trans* signaling was triggered by pheromone (i.e., within 30-50% of wild-type counterparts). There was no signaling output if either mutant lacked the CTM domain, indicating that both mutants must be membrane-tethered for *trans* signaling to occur (Figures 3B). Notably, Gβγ played no role here, because the strains lacked the gene for the Gβ subunit (Ste4). Furthermore, the results were unchanged when the PM and RING-H2 domains were removed by deleting the Ste5 N-terminus (ΔN, Figure 3B); thus, when the role of these domains in membrane localization were bypassed, so too were their roles *in trans* signaling. We conclude that interaction of Gβγ with the RING-H2 domain is not strictly required for *trans* signaling, and that their normal requirement can be explained by their role in membrane recruitment.

**FIGURE 3:**
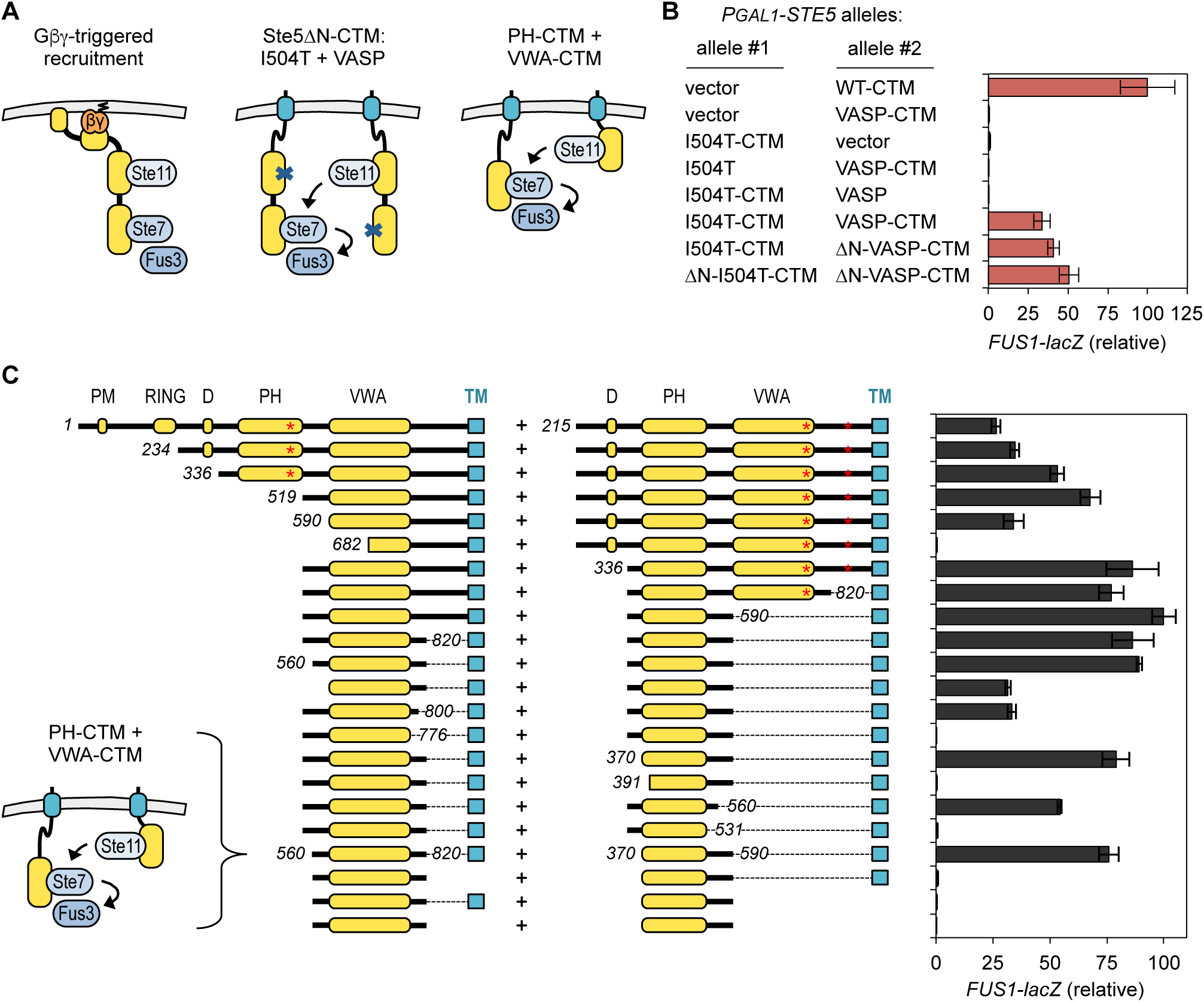
Membrane-tethered Ste5 allows *trans* signaling and definition of minimal domains. (A) Cartoons of normal Ste5 membrane recruitment (*left*) and *trans* signaling by membrane-tethered forms of Ste5 (*center, right*), as assayed in the next panels. For simplicity of illustration, the C-terminal transmembrane (CTM) domain is shown in a generic position. (B) *Trans* signaling by Ste5-CTM derivatives bypasses requirement for N-terminal PM and RING-H2 domains. *FUS1-lacZ* induction (mean ± SD; n = 4) was measured in *ste4Δ ste5Δ* cells (PPY886) coexpressing mutant variants of Ste5 or Ste5ΔN (residues 215-917), with or without a CTM domain. *P*_*GAL1*_-driven constructs were induced with galactose (3 hr). (C) Minimal domains for *trans* signaling. Experiments were performed as in panel B, using various truncations of Ste5-CTM. Red asterisks show positions of I504T and VASP mutations; numbers denote positions of new truncation endpoints. Bars, mean ± SD (n = 3). Also see Figure S2B-D.

### Definition of minimal domains for *trans* signaling

Next, we used the Ste5-CTM fusions to determine the minimal domains of Ste5 required for *trans* signaling. We reasoned that artificial membrane tethering should allow removal of all sequences needed only for membrane localization, and thus any remaining requirements would define regions critical for signaling between kinases. In addition, because the two complementing molecules perform distinct roles, we expected that each partner could be reduced to the minimum sequence needed to perform its individual role, free of other constraints that might be imposed when a single polypeptide performs all roles.

Indeed, using a series of truncations, we found that each partner could be trimmed to a region comprising a single structural domain (Figures 3C, S2A). The partner that provides the Ste11-binding role could be trimmed to a fragment corresponding to the PH domain (residues 370-590), and its counterpart could be trimmed to a fragment corresponding to the VWA domain (residues 560-820). In each case, some further truncation was tolerated but led to reduced function (e.g., when the PH domain C-terminus was trimmed to 560, and when the VWA domain was trimmed at its N-terminus to 590 or at its C-terminus to 800). These partial reductions might reflect a simple need for linker sequence, such as to allow sufficient separation from the membrane or the CTM domain; indeed, using the original complementation assay between Ste5-I504T and Ste5-VASP (Figure S2B), the C-terminus of the PH domain could be truncated even further (to residue 531). Ultimately, robust signaling could be achieved by combining the two minimal fragments encoding the PH and VWA domains (Figure 3B). Further controls showed that both fragments must be membrane-tethered (Figure 3B); this finding is notable because the isolated VWA domain should be relieved of auto-inhibition [16], and yet its ability to transmit signal still required membrane localization. Other controls using monomeric YFP and mCherry to replace an N-terminal GFP moiety showed that the signaling results did not rely on potential dimerization tendencies of GFP (Figure S2C). Collectively, the results show that *trans* signaling can be induced by simultaneously targeting to the membrane two minimal signaling domains of Ste5 – the PH and VWA domains – that provide two distinct functions. In this context, other sequences were dispensable.

### Stimulus-induced regulation of individual functions

The preceding results suggested to us that it should be possible to split the functions of Ste5 in two, and then have only one function controlled by the pheromone stimulus. To address this possibility, we used the membrane-tethered domains defined in the previous section (PH-CTM or VWA-CTM), and then asked if pheromone could still control pathway output by regulating the remainder of Ste5 (i.e., lacking the excised domain). For these experiments, when removing the Ste5 PH domain we replaced it with a heterologous PH domain from mammalian PLCδ (i.e., Ste5ΔPH::PLC), to compensate for any defects in membrane localization.

Initially, we confirmed that the mutants lacking individual domains (Ste5-ΔVWA and Ste5-ΔPH::PLC) were able to complement each other, and that each partner required its localization sequences to remain intact (Figure 4A,B). Then, we tested each mutant for complementation by a membrane-tethered version of the excised domain (Figure 4C,D), and found that pathway output was still regulated by pheromone in each case (i.e., Ste5ΔVWA + VWA-CTM or Ste5ΔPH::PLC + PH-CTM). Because in either context only one partner contained a Gβγ-binding domain, the results imply that pheromone controlled the ability of that partner to functionally engage with its membrane-localized counterpart. Furthermore, each pheromone-regulated partner still required the remaining localization sequences (PM and RING) (Figure 4B). Thus, separation of Ste5 into distinct functional portions allows pathway regulation by pheromone and Gβγ to be conferred by either portion individually, without having all portions regulated simultaneously. Given that the PH and VWA domains can function on different polypeptides, we also asked if their order within the same polypeptide could be interchanged. Indeed, the resulting domain-swapped form of Ste5 functioned similarly to wild-type Ste5 (Figure S3). This finding further illustrates the functional modularity of Ste5, and it places constraints on how any conformational changes that might be triggered by Gβγ binding could be propagated throughout the rest of the polypeptide. Overall, these results reveal flexibility in the point of regulatory input, and they provide a setting in which it is possible to dissect distinct stimulus-regulated events.

**FIGURE 4:**
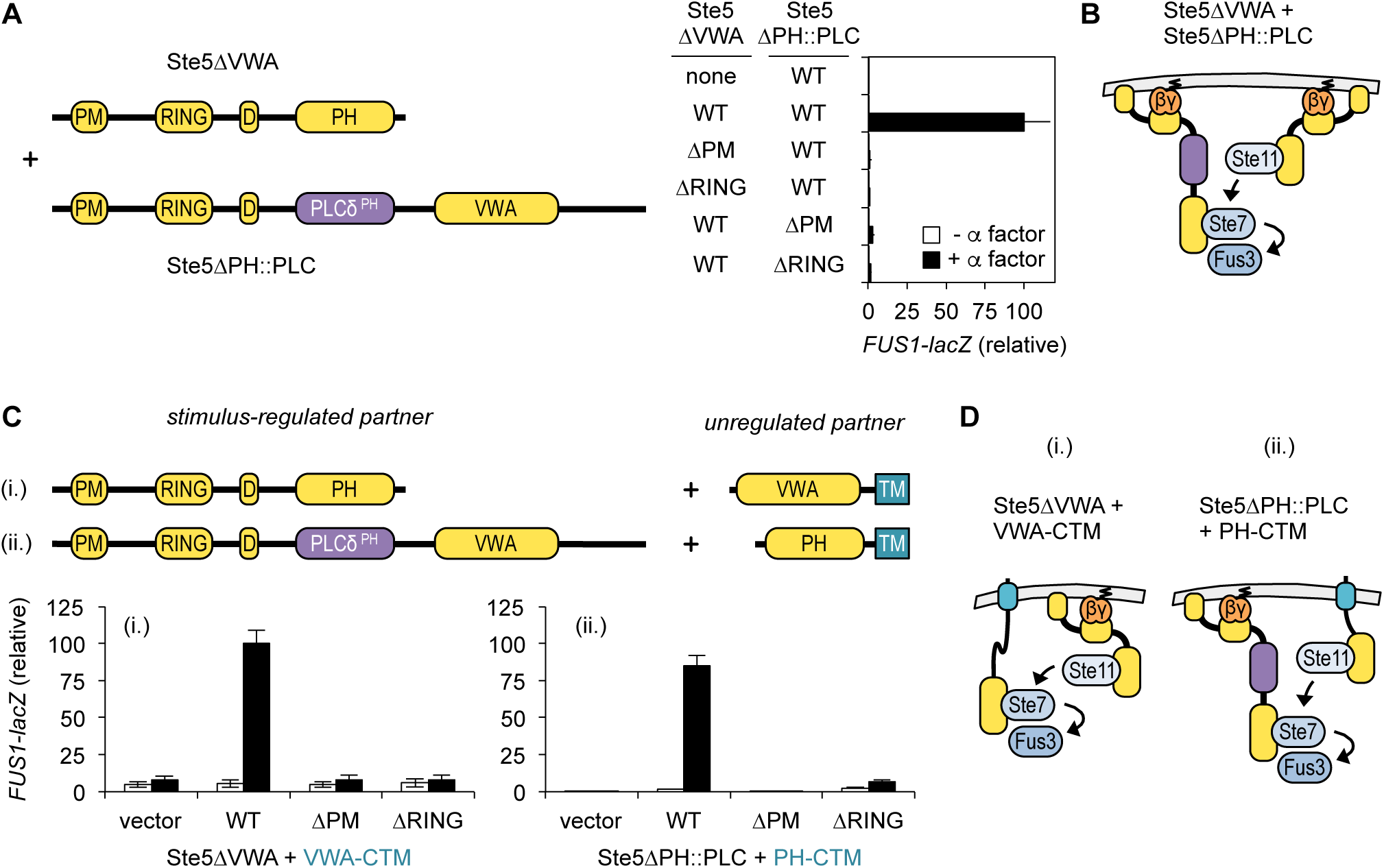
Stimulus-mediated regulation of individual Ste5 functions. (A) Coexpression of Ste5ΔVWA and Ste5ΔPH::PLC (diagrammed at left) yields *trans* signaling, and both require intact PM and RING-H2 domains. *FUS1-lacZ* induction (mean ± SD; n = 4) was assayed ± α factor (5 µM, 2 hr.). Strain PPY2032 harbored HA_3-_ and myc_13_-tagged variants of Ste5ΔVWA and Ste5ΔPH::PLC, respectively. (B) Cartoon depiction of the signaling scenario in panel A. (C) Combination of a stimulus-regulated partner (Ste5ΔVWA or Ste5ΔPH::PLC) with an unregulated, membrane-tethered partner (VWA-CTM or PH-CTM) yields stimulus-regulated signaling. *FUS1-lacZ* induction (mean ± SD; n = 4) was assayed after induction with galactose ± α factor (5 µM, 3 hr). Strain: PPY858. Also see Figure S3. (D) Cartoon depictions of the signaling scenarios in panel C.

### *Trans* signaling by a membrane-localized, scaffold-free kinase

We next explored whether directly tethering one of the pathway kinases to the membrane could eliminate the need for a scaffold protein to localize it. For this we used a membrane-tethered form of Ste11 (Ste11-Cpr), which contains a C-terminal “CCaaX” motif that gets modified with lipophilic groups [10]. In the *trans*-signaling assay, we asked if this Ste11-Cpr fusion could substitute for Ste5-VASP (the partner that normally provides the Ste11-binding role) and thus complement Ste5-I504T (Figure 5A). Indeed, in a *ste5-I504T* strain, expression of Ste11-Cpr on its own was not sufficient to activate signaling, but it could do so when pheromone was added. This response required Ste11 to be membrane-tethered, as it was not observed if the key Cys residues in the CCaaX motif were mutated (Cpr-SS). Thus, in this context *trans* signaling occurred between membrane-bound Ste11 and scaffold-bound Ste7 (Figure 5A, right). Furthermore, we found that Ste11-Cpr could also activate signaling when co-expressed with membrane-tethered forms of the Ste5 VWA domain (Figures 5B, S4A). Both observations emphasize that *trans* signaling is not inherently dependent on specific inter-scaffold contacts.

**FIGURE 5:**
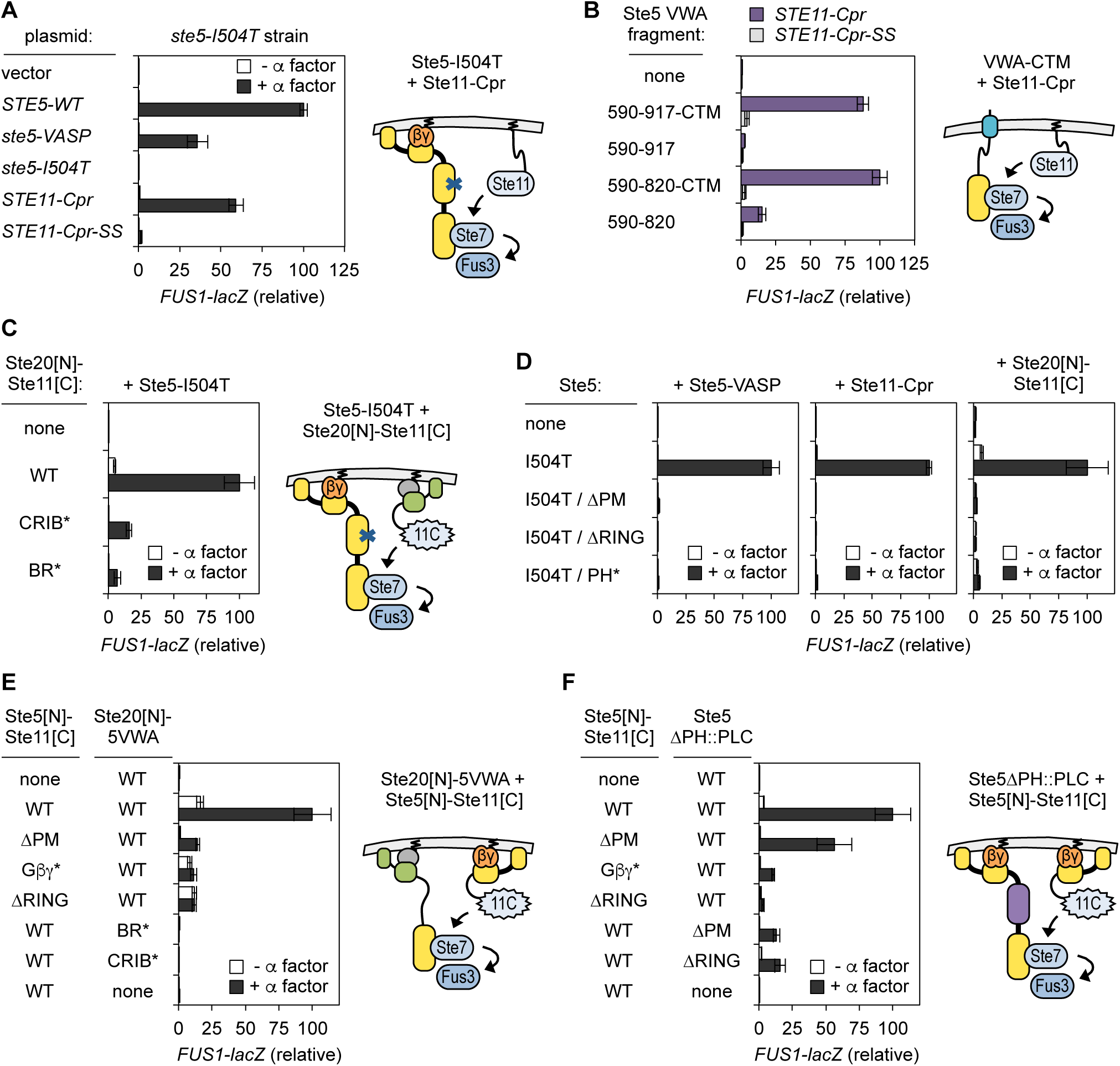
*Trans* signaling by membrane-tethered kinases and alternate recruitment methods. (A) Complementation of the Ste5-I504T signaling defect by membrane-tethered Ste11 (Ste11-Cpr). *FUS1-lacZ* induction (mean ± SD; n = 4) was assayed in *ste5-I504T* cells (PPY1968) harboring *STE5-myc*_*13*_ or *P*_*GAL1*_*-STE11-Cpr* plasmids, after induction with galactose ± α factor (5 µM, 3 hr). (B) *FUS1-lacZ* induction (mean ± SD; n = 4) in cells (PPY2252) coexpressing the indicated *P*_*GAL1*_*-STE5-CTM* and *P*_*GAL1*_*-STE11-Cpr* constructs, induced at moderate strength by using 10 nM β-estradiol (90 min). See also Figure S4A. (C) Ste5-I504T is complemented by Ste20[N]-Ste11[C], where the active Ste11 kinase domain is localized by the Ste20 N-terminus. *FUS1-lacZ* (mean ± SD; n = 4) was assayed ± α factor (5 µM, 2 hr). CRIB* and BR* denote mutations in Ste20 sequences that bind Cdc42 and plasma membrane, respectively [50, 51]. Strain: PPY2032. (D) Localization sequences are required for Ste5-I504T to complement Ste5-VASP, Ste11-Cpr, or Ste20[N]-Ste11[C]. *FUS1-lacZ* induction (mean ± SD) was assayed ± α factor (5 µM, 2 hr; n = 4; PPY2032), or for Ste11-Cpr after induction with galactose ± α factor (5 µM, 3 hr; n = 3; PPY858). Also see Figure S4A. (E) *FUS1-lacZ* induction (mean ± SD; n = 6) in *ste5Δ* cells (PPY2032) harboring plasmid-borne Ste5[N]-Ste11[C] and Ste20[N]-Ste5[VWA] variants, treated ± α factor (5 µM, 2 hr). (F) *FUS1-lacZ* induction (mean ± SD; n = 6) in *ste5Δ* cells (PPY2032) harboring plasmid-borne Ste5[N]-Ste11[C] and Ste5ΔPH::PLC variants, treated ± α factor (5 µM, 2 hr).

We investigated two further methods to drive signaling by kinase co-localization. In one case we used a construct, Ste20[N]-Ste11[C], in which the C-terminal kinase domain from Ste11 is fused to localization sequences from the Ste20 N-terminus (Figure 5C, right). This fusion was previously shown to increase signaling in wild-type and *ste11*Δ cells [23, 24]. Here, we found that it also could mediate *trans*-signaling with Ste5-I504T (Figure 5C), which was disrupted by mutations in the localization sequences from Ste20 that bind Cdc42 (CRIB*) or the plasma membrane (BR*). Because the Ste11[C] fragment includes only the catalytic domain of Ste11, it is expected to be constitutively active due to being freed from its inhibitory N-terminus [12]; therefore, the finding that its *trans*-signaling activity still required membrane localization implies that signaling was stimulated by co-localizing the active kinase with its substrate. Its partner molecule, Ste5-I504T, also required intact localization sequences, and this was true regardless of which specific molecule it was paired with (Figures 5D, S4B).

In another case, we used the Ste20 N-terminus to localize the Ste5 VWA domain (Ste20[N]-5VWA) (Figure 5E, right), and then combined this chimera with Ste5[N]-Ste11[C] [23], in which the Ste11 catalytic domain is fused to the N-terminus of Ste5 (containing the PM and RING-H2 domains). This combination also yielded *trans* signaling, which again relied on localization sequences in each partner (Figure 5E). This result is significant because the two chimeric proteins should constitutively achieve two distinct activation steps that are normally controlled by pheromone (i.e., de-repression of both the Ste11 kinase domain and the Ste5 VWA domain), and yet signal output was still regulated by pheromone and localization. There was a notable increase in basal signaling (∼16% of maximum), suggesting that these deregulations impose a cost of promiscuous signaling and reduced dynamic range. Basal signaling was lower (< 4% of maximum) in a related context (Figure 5F) in which membrane localization of the Ste5 VWA domain was not constitutive but instead remained pheromone-induced (using Ste5ΔPH::PLC). Curiously, in this arrangement there was greater tolerance for deleting the PM domain from the Ste5[N]-Ste11[C] chimera; the reasons for this are not known, but previous work indicates that the PM domain helps increase net affinity for membrane-bound Gβγ and can become less essential when Ste5 levels are elevated [10], and so perhaps this reduced affinity can also become tolerated when other critical pathway steps are enhanced or accelerated (e.g., due to release from inhibition). Altogether, these findings reveal a remarkable variety of arrangements in which kinase colocalization allows *trans* signaling while still retaining stimulus-mediated regulation.

### No evidence for MAPK activation *in trans*

Finally, we asked if the Ste7 → Fus3 reaction might also be able to occur *in trans*. To address this question, we co-expressed two Ste7 mutants (Figure S4C): one that lacks kinase activity but retains Fus3 binding sequences (Ste7-R220; [25]) and one that retains kinase activity but cannot bind Fus3 (Ste7-ND; [26]). We observed no complementation between these forms. While less conclusive than a positive result, this finding might imply that Ste5 can only promote the Ste7 → Fus3 reaction in *cis*. We sought to probe this issue further using mutations in the Ste5 VWA domain, which has two distinct functional surfaces (Figure S4D): one that binds Ste7 and one that induces Fus3 to become a good substrate for Ste7 [22]. Unfortunately, while mutations in either region can severely disrupt the reaction in vitro (by 100-1000x; [22]), we found their impact on signaling in vivo to be surprisingly mild (< 2x) (Figure S4E), and hence unsuitable for complementation tests. Curiously, stronger defects emerged (Figure S4F) when signaling was initiated late in the pathway by a pseudo-active form of Ste7 (Ste7-EE), which was the only input source used in the prior in vitro studies [22]. This context dependence was unexpected, and it likely holds mechanistic implications that will be of interest to future studies.

## DISCUSSION

Our findings illuminate how signal transmission can be stimulated by colocalization of signaling proteins to the plasma membrane. In particular, we report that membrane localization of Ste5 promotes signaling *in trans* between kinases that are bound to separate scaffold molecules. This conclusion is supported by the findings that *trans* signaling requires all localization sequences in Ste5, that direct membrane tethering of either Ste5 or a pathway kinase can suffice to promote *trans* signaling, and that *trans* signaling can be activated by a diverse array of methods that achieve kinase co-localization. Moreover, the *trans* signaling reaction allowed us to divide Ste5 into distinct functional domains, and to achieve normal regulation of signaling when either one was stimulated to colocalize with its membrane-tethered counterpart. The overall findings suggest that the scaffold mediates membrane colocalization of pathway kinases, which can enhance their signaling interactions by concentrating them in a reduced subcellular volume (Figure 6). This property can relax assembly constraints for scaffold proteins and increase regulatory flexibility, which has broad relevance to the function and evolution of signaling pathways.

**FIGURE 6:**
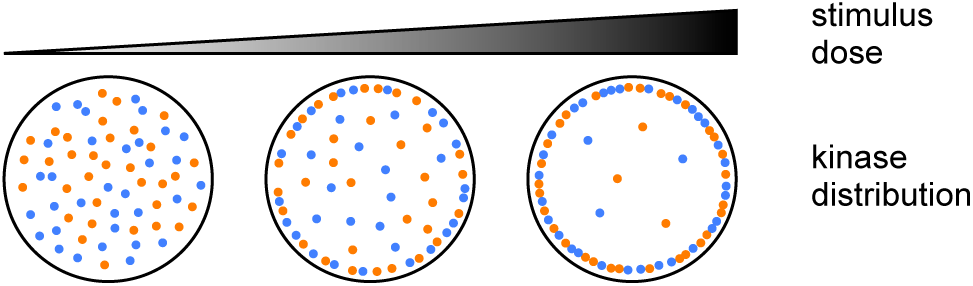
General model for concentrating kinases by membrane colocalization. As the stimulus dose increases, an increasing fraction of molecules is redistributed to a reduced volume of cytoplasm adjacent to the plasma membrane. The increased concentration can favor protein-protein interactions and catalytic reactions among signaling proteins. Results in this study suggest that a scaffold protein can mediate this process by serving as a membrane-recruited binding sink for distinct kinases that signal to each other without being bound to the same scaffold molecule.

It is relatively straightforward to envision how membrane recruitment of a cytoplasmic protein can promote its interaction with a membrane-localized partner (e.g., as in the initial Ste20 → Ste11 step). Yet membrane recruitment can also promote interactions between two cytoplasmic proteins, by strongly increasing their local concentrations [2]. In theory, translocation of proteins to a reduced volume of cytoplasm adjacent to the plasma membrane can raise local concentrations by 100-to 1000-fold [1, 27], which can increase the number of complexes between two proteins when even a moderate fraction of each (e.g., 10%) is colocalized. Scaffold proteins are themselves expected to enforce mutual proximity between their bound partners, but membrane colocalization could still be beneficial if scaffold molecules are not fully occupied. This is necessarily true for the mutant scaffolds used in this study, yet even in wild-type cells only a minority of Ste5 molecules (∼ 10-25%) are bound to any given kinase [7, 28]. Thus, because membrane localization allows Ste5 to enhance signal propagation without any individual scaffold molecules being fully occupied with kinases, it suggests a general scenario in which scaffolds need not assemble individual multiprotein complexes, but instead can promote signaling by serving as binding sinks that concentrate reactants in specific subcellular regions. This scenario loosens functional constraints on the scaffold and hence fits with models in which signaling is governed by heterogeneous protein “ensembles” [29], rather than by homogeneous, unitary complexes. This alternative view can help explain effects of membrane localization on signal output in previous studies [13, 14], as well as the remarkable ease with which signaling networks can be reconfigured [24].

In principle, even stronger concentration effects could be achieved by recruiting proteins into the smaller volume of a specialized microdomain, for which there is precedent in metazoan MAPK pathways and other signaling systems [30, 31]. It is unclear if such domains exist in the yeast pheromone pathway, although in some reports Ste5 has appeared localized to membrane puncta [8, 11, 32], the nature of which is currently unknown. Signaling reactions can also be promoted by the opposite of colocalization: namely, by excluding enzymes that catalyze the reverse reactions, such as phosphatases. In the pheromone pathway, phosphatases that inactivate the MAPK are not co-recruited with it to the membrane, such that the active MAPK is partitioned from its antagonists [7]. So far, however, comparable information is not available for phosphatases that reverse the upstream reactions.

It remains unknown whether the Ste11 → Ste7 reaction in normal cells occurs more frequently *in trans* or *in cis*. This is a challenging question to probe experimentally, because while it is easy to devise a context in which signaling must occur *in trans* (e.g., using the mutants in this study), it is very difficult to devise one in which signaling must occur *in cis*; hence, at present we cannot directly compare their efficiencies or even verify that the *cis* reaction occurs. In our assays, the reduction in pathway output when *trans* signaling was required (compared to cells with WT Ste5) might roughly estimate the fraction of signaling that normally occurs *in cis*, although more complex scenarios cannot be excluded. There is no evidence that a steric constraint forbids the *cis* reaction, and instead current data favor a flexible linkage between the kinase-binding domains of Ste5. Thus, we suspect that the relative frequency of *cis* versus *trans* reactions is most likely dictated by the kinase occupancy of Ste5 and the local concentration achieved by membrane recruitment. This issue remains a worthwhile subject for future studies, including those involving computational approaches.

*Trans* signaling behavior was originally attributed to Ste5 dimerization, but our findings suggest that it can be explained more readily by membrane colocalization of multiple Ste5 molecules. Indeed, the need for all localization sequences in Ste5 indicates that the earlier observed role for the RING-H2 domain does not imply a unique dimerization function but instead exemplifies a more general need for membrane recruitment. To date there are no examples of Ste5 mutants whose signaling defects can be clearly attributed to a dimerization defect. Biophysical methods show that full-length Ste5 is predominantly monomeric in intact cells [7, 28], although co-immunoprecipitation can be detected in cell extracts [11, 33]. Two-hybrid assays can detect self-interaction for a Ste5 fragment (residues 25-587), but this does not require the RING-H2 domain ([17]; M.J.W. and P.M.P., unpublished observations). The isolated RING-H2 domain does not show appreciable self-interaction when fully intact [10, 34], but it does when perturbed by partial truncation [10] or mutation of zinc-chelating cysteines (M.J.W. and P.M.P., unpublished observations). It is also relevant that dimerization might be expected to allow Ste5 localization defects to be compensated by a localization-competent partner, but instead we found that the kinase-binding mutants could not complement any of the localization mutants. With the benefit of hindsight and further advances in the field, we suggest the most parsimonious interpretation is that *trans* signaling does not necessarily require Ste5-Ste5 contacts but instead results from the colocalization of multiple Ste5 molecules at the membrane. It remains possible, of course, that Ste5-Ste5 contacts could make such reactions even more efficient by further increasing local concentration beyond that achieved by colocalization alone.

MAPK pathways exist in all eukaryotes and have conserved functions. Despite its essential role in pheromone response, Ste5 is not conserved as strongly as its pathway kinases [35], and hence it likely emerged as an addition to a pre-existing pathway. Indeed, membrane localization of the homologous MAPK cascade in filamentous fungi depends on a scaffold protein (HAM-5) that is unrelated to Ste5 [36, 37]. Moreover, scaffold proteins for MAPK pathways in metazoans (e.g., KSR, JIP-1, etc.) are not related to their fungal analogs or to each other [38], suggesting that this functional category has evolved independently multiple times. The ability to acquire a new scaffolding function might be assisted by the properties studied here. The stimulatory effects of membrane colocalization provide substantial flexibility in the mechanism of localization, and allow different steps to be driven by different scaffold molecules or even different proteins. Thus, localization could provide a simple way of gaining a new regulatory input, which could then foster evolution of more complex features such as specific binding interactions and allosteric changes [2, 35]. These notions complement those arising from other studies on the yeast pheromone pathway, in which domain shuffling among pathway components can generate novel signaling behaviors and reveal the tolerance of the network to reconfiguration, both of which can facilitate evolution [23, 24].

Collectively, the findings reported here and in prior studies [5, 12, 16] suggest that membrane recruitment separately promotes all three kinase activation steps in the pheromone pathway; i.e., activation of Ste11 by membrane-localized Ste20, *trans*-activation of Ste7 by Ste11, and de-repression of the Ste5 VWA domain to allow activation of Fus3 by Ste7. Moreover, pheromone can still control pathway output when some of these steps are activated constitutively; i.e., when Ste11 is pre-activated by mutation [13, 14], or when the Ste5 VWA domain is freed from its *cis*-inhibitory PH domain (this study), or even both (this study). Thus, this pathway exhibits elements of both fine-tuning and coarse functional flexibility. That is, the ability to control multiple steps simultaneously could help increase the dynamic range of response and reduce basal signaling, while also imparting tolerance of suboptimal parameters for individual reactions. Moreover, the ability to control pathway output even when some steps are constitutively active could foster the evolution of new pathways or new pathway stimuli, by allowing diversity in the regulated step [39, 40]. By analogy, design of synthetic signaling pathways [41, 42] is likely to benefit from an evolutionary process in which simple initial circuits are gradually refined by incorporating additional control points.

## METHODS

### Yeast Strains and Growth Conditions

Standard procedures were used for growth and genetic manipulation of yeast [43, 44]. Yeast strains were in W303 or S288C (derived from YPH499) backgrounds. Cells were grown at 30°C in yeast extract/peptone medium with 2% glucose (YPD), or in synthetic (SC) medium (lacking specific nutrients appropriate to select for plasmids) with 2% glucose or raffinose. Strains and plasmids are listed in Tables S1 and S2, respectively. Mutations at the native *ste5* genomic locus (I504T and VASP) were introduced by a two-step allele replacement method [43] using plasmids pPP2870 and pPP2871.

### Mating and Pheromone Response Assays

Patch mating assays were performed between test strains (*MAT***a**) and a partner strain (PT2α) using methods described previously [45]. To measure transcriptional responses, cells harboring an integrated *FUS1-lacZ* reporter were treated with 5 µM α factor for 2 hr. For signaling induced by *P*_*GAL1*_-regulated genes, cells growing in SC medium with raffinose were supplemented with 2% galactose, and incubated for 3 hr (with or without 5 µM α factor); in some experiments, *P*_*GAL1*_-regulated genes were induced at submaximal strength by using a hybrid transcription factor (Gal4^DBD^-hER-VP16, or “GEV”) whose activity was controlled by the hormone β-estradiol [14]. Afterward, cells were collected and assayed for β-galactosidase activity by colorimetric assay as described previously [21]. To measure MAPK phosphorylation, cells were treated ± α factor (5 µM, 15 min.) or, where applicable, treated first with galactose for 90 min. and then incubated ± α factor (5 µM, 30 min.). Afterward, 2-mL samples were harvested by centrifugation, then cell pellets were immediately frozen in liquid nitrogen and stored at −80°C.

### Cell Extracts and Immunoblotting

Whole cell extracts were prepared by lysis in trichloroacetic acid as described previously [46], using frozen cell pellets from 2 mL cultures. Protein concentrations were measured by bicinchoninic acid (BCA) assay (Pierce #23225), and equal amounts (10 µg) were loaded per lane. Proteins were resolved by SDS-PAGE and transferred to polyvinylidene fluoride (PVDF) in a submerged tank. Primary antibodies were rabbit anti-phospho-p44/42 (1:1000, Cell Signaling Technology #9101), rabbit anti-myc (1:200 Santa Cruz Biotechnologies #sc-789), rabbit anti-G6PDH (1:100000, Sigma #A9521), or mouse anti-HA (1:1000, Covance #MMS101R). HRP-conjugated secondary antibodies were goat anti-rabbit (1:3000, Jackson ImmunoResearch #111-035-144) or goat anti-mouse (1:3000, BioRad #170-6516). Enhanced chemilluminescent detection used a BioRad Clarity substrate (#170-5060). Exposures were captured on x-ray film.

## SUPPLEMENTAL INFORMATION

Supplemental Information includes four figures and two tables.

### ACKNOWLEDGEMENTS

This work was supported by a grant from the NIH (R01 GM057769) to P.M.P. We thank Beverly Errede, Wendell Lim, and Jeremy Thorner for sharing yeast plasmids, plus Alejandro Colman-Lerner and Dan McCollum for feedback on the manuscript.

## AUTHOR CONTRIBUTIONS

R.E.L. and M.J.W. performed the experiments; P.M.P. designed the project, analyzed the data, and wrote the paper.

