## Supplemental Information for "Ste5 Membrane Localization Allows MAPK Pathway Signaling *in trans* Between Kinases on Separate Scaffold Molecules"

for

**TABLE S1: Yeast strains used in this study.**

| Strain background* | Name | Relevant genotype | Source |
|---|---|---|---|
| a | PPY1955 | *MAT***a** *FUS1::FUS1-lacZ::LYS2* | this study |
| a | PPY1974 | *MAT***a** *FUS1::FUS1-lacZ::LYS2 ste5-VASP* | this study |
| a | PPY1975 | *MAT***a** *FUS1::FUS1-lacZ::LYS2 ste5-I504T* | this study |
| a | PPY2032 | *MAT***a** *FUS1::FUS1-lacZ::LYS2 ste5::ADE2* | this study |
| b | PPY640 | *MAT***a** *FUS1::FUS1-lacZ::LEU2* | [S1] |
| b | PPY858 | *MAT***a** *FUS1::FUS1-lacZ::LEU2 ste5::ADE2* | [S1] |
| b | PPY886 | *MAT***a** *FUS1::FUS1-lacZ::LEU2 ste4::*$ura3^{FOA}$ *ste5::ADE2* | [S1] |
| b | PPY891 | *MAT***a** *FUS1::FUS1-lacZ::LEU2 ste7::ADE2* | [S1] |
| b | PPY1669 | *MAT***a** *FUS1::FUS1-lacZ::LEU2 ste5::ADE2 kss1::*$ura3^{FOA}$ | this study |
| b | PPY1967 | *MAT***a** *FUS1::FUS1-lacZ::LEU2 ste5-VASP* | this study |
| b | PPY1968 | *MAT***a** *FUS1::FUS1-lacZ::LEU2 ste5-I504T* | this study |
| b | PPY2252 | *MAT***a** *FUS1::FUS1-lacZ::LEU2 ste5::ADE2 TRP1::*$P_{ADH1}$*-GEV* | this study |
| c | PT2$\alpha$ | *MAT*$\alpha$ *hom3 ilv1 can1* | [S1] |

* (a) S288C [*ade2-101 his3-*$\Delta$*200 leu2-*$\Delta$*1 lys2-801 trp1-*$\Delta$*63 ura3-52*] (b) W303 [*ade2-1 his3-11,15 leu2-3,112 trp1-1 ura3-1 can1*]; (c) other.

**TABLE S2: Plasmids used in this study.**

| Name | Description * | Source |
|---|---|---|
| pFD-STE11-Asp3 | CEN ARS URA3 STE11-Asp3 | [S2] |
| pL38-WT | CEN ARS HIS3 $P_{GAL1}$-STE4 | [S3] |
| pNC589 | CEN ARS TRP1 $P_{CYC1}$-ste7(EE)-myc | [S4] |
| pPP511 | CEN ARS TRP1 $P_{GAL1}$-GFP-STE5-CTM | [S1] |
| pPP1532 | CEN ARS HIS3 $P_{GAL1}$-STE11-Cpr | [S5] |
| pPP1815 | CEN ARS URA3 ste7(K220R)-$GFP_{x3}$ $T_{CYC1}$ | this study |
| pPP1969 | CEN ARS URA3 $P_{STE5}$-STE5-$myc_{13}$ $T_{CYC1}$ | [S5] |
| pPP2019 | CEN ARS URA3 $P_{GAL1}$-GFP vector | [S5] |
| pPP2104 | CEN ARS URA3 $P_{STE5}$-ste5(ΔPM)-$myc_{13}$ $T_{CYC1}$ | [S5] |
| pPP2211 | CEN ARS HIS3 $P_{GAL1}$-STE11-Cpr-SS | [S5] |
| pPP2544 | CEN ARS URA3 $P_{STE5}$-ste5(ΔRING)-$myc_{13}$ $T_{CYC1}$ | [S6] |
| pPP2857 | CEN ARS TRP1 $P_{STE5}$-STE5-$HA_3$ $T_{CYC1}$ | [S7] |
| pPP2861 | CEN ARS URA3 $P_{STE5}$-ste5(VASP)-$myc_{13}$ $T_{CYC1}$ | [S6] |
| pPP2862 | CEN ARS URA3 $P_{STE5}$-ste5(I504T)-$myc_{13}$ $T_{CYC1}$ | [S6] |
| pPP2863 | CEN ARS TRP1 $P_{STE5}$-ste5(VASP)-$HA_3$ $T_{CYC1}$ | [S7] |
| pPP2870 | integrating URA3 ste5(VASP) | this study |
| pPP2871 | integrating URA3 ste5(I504T) | this study |
| pPP2879 | CEN ARS URA3 $P_{GAL1}$-GFP-ste5(Δ1-214/I504T)-CTM | this study |
| pPP2902 | CEN ARS TRP1 $P_{GAL1}$-GFP-ste5(Δ1-214/VASP)-CTM | this study |
| pPP2907 | CEN ARS URA3 $P_{GAL1}$-GFP-ste5-I504T | this study |
| pPP2908 | CEN ARS TRP1 $P_{GAL1}$-GFP-ste5(VASP) | this study |
| pPP3015 | CEN ARS TRP1 $P_{STE5}$-ste5(VASP/ΔPM)-$HA_3$ $T_{CYC1}$ | this study |
| pPP3016 | CEN ARS TRP1 $P_{STE5}$-ste5(VASP/ΔRING)-$HA_3$ $T_{CYC1}$ | this study |
| pPP3017 | CEN ARS TRP1 $P_{STE5}$-ste5(VASP/Gβγ*)-$HA_3$ $T_{CYC1}$ | this study |
| pPP3018 | CEN ARS TRP1 $P_{STE5}$-ste5(VASP/ΔPM::PLCδ$^{PH}$)-$HA_3$ $T_{CYC1}$ | this study |
| pPP3019 | CEN ARS TRP1 $P_{STE5}$-ste5(VASP/PH*)-$HA_3$ $T_{CYC1}$ | this study |
| pPP3020 | CEN ARS TRP1 $P_{STE5}$-ste5(VASP/ΔPM::2xPLCδ$^{PH}$)-$HA_3$ $T_{CYC1}$ | this study |
| pPP3048 | CEN ARS URA3 $P_{STE5}$-ste5(PH*)-$myc_{13}$ $T_{CYC1}$ | this study |
| pPP3128 | CEN ARS URA3 $P_{STE5}$-ste5(Gβγ*)-$myc_{13}$ $T_{CYC1}$ | this study |
| pPP3130 | CEN ARS URA3 $P_{GAL1}$-GFP-ste5(I504T)-CTM | this study |
| pPP3131 | CEN ARS URA3 $P_{GAL1}$-GFP-ste5(Δ1-233/I504T)-CTM | this study |
| pPP3132 | CEN ARS URA3 $P_{GAL1}$-GFP-ste5(Δ1-335/I504T)-CTM | this study |
| pPP3134 | CEN ARS URA3 $P_{GAL1}$-GFP-ste5(Δ1-518)-CTM | this study |
| pPP3135 | CEN ARS URA3 $P_{GAL1}$-GFP-ste5(Δ1-589)-CTM | this study |
| pPP3136 | CEN ARS URA3 $P_{GAL1}$-GFP-ste5(Δ1-681)-CTM | this study |
| pPP3141 | CEN ARS URA3 $P_{GAL1}$-GFP-ste5(519-820)-CTM | this study |
| pPP3142 | CEN ARS URA3 $P_{GAL1}$-GFP-ste5(590-820)-CTM | this study |
| pPP3143 | CEN ARS TRP1 $P_{GAL1}$-GFP-ste5(336-917/V763A S861P)-CTM | this study |
| pPP3144 | CEN ARS TRP1 $P_{GAL1}$-GFP-ste5(336-820/V763A)-CTM | this study |
| pPP3145 | CEN ARS TRP1 $P_{GAL1}$-GFP-ste5(336-590)-CTM | this study |
| pPP3163 | CEN ARS URA3 $P_{GAL1}$-GFP-ste5(560-820)-CTM | this study |
| pPP3164 | CEN ARS URA3 $P_{GAL1}$-GFP-ste5(519-800)-CTM | this study |
| pPP3165 | CEN ARS URA3 $P_{GAL1}$-GFP-ste5(519-776)-CTM | this study |
| pPP3166 | CEN ARS TRP1 $P_{GAL1}$-GFP-ste5(370-590)-CTM | this study |
| pPP3167 | CEN ARS TRP1 $P_{GAL1}$-GFP-ste5(391-590)-CTM | this study |
| pPP3168 | CEN ARS TRP1 $P_{GAL1}$-GFP-ste5(336-560)-CTM | this study |
| pPP3169 | CEN ARS TRP1 $P_{GAL1}$-GFP-ste5(336-531)-CTM | this study |
| pPP3176 | CEN ARS TRP1 $P_{STE5}$-ste5(1-590)-$HA_3$ $T_{CYC1}$ | this study |
| pPP3210 | CEN ARS URA3 $P_{GAL1}$-GFP-ste5(560-820) | this study |
| pPP3211 | CEN ARS TRP1 $P_{GAL1}$-GFP-ste5(370-590) | this study |
| pPP3226 | CEN ARS TRP1 $P_{STE5}$-ste5(1-560)-$HA_3$ $T_{CYC1}$ | this study |
| pPP3229 | CEN ARS URA3 $P_{STE5}$-ste5(Δ364-532)-$myc_{13}$ $T_{CYC1}$ | this study |
| pPP3233 | CEN ARS URA3 $P_{STE5}$-ste5(Δ364-532::PLCδ$^{PH}$)-$myc_{13}$ $T_{CYC1}$ | this study |
| pPP3321 | CEN ARS URA3 $P_{STE5}$-ste5(coa1)-$myc_{13}$ $T_{CYC1}$ | this study |
| pPP3322 | CEN ARS URA3 $P_{STE5}$-ste5(coa2)-$myc_{13}$ $T_{CYC1}$ | this study |
| pPP3323 | CEN ARS URA3 $P_{STE5}$-ste5(coa3)-$myc_{13}$ $T_{CYC1}$ | this study |
| pPP3324 | CEN ARS URA3 $P_{STE5}$-ste5(Δ7BR)-$myc_{13}$ $T_{CYC1}$ | this study |
| pPP3331 | CEN ARS URA3 $P_{STE5}$-ste5(1-560)-$myc_{13}$ $T_{CYC1}$ | this study |
| pPP3334 | CEN ARS URA3 $P_{STE5}$-ste5([Δ364-532]-[370-590])-$myc_{13}$ $T_{CYC1}$ | this study |
| pPP3444 | CEN ARS TRP1 $P_{GAL1}$-mYFP-ste5(Δ1-214/VASP)-CTM | this study |

| | | |
|---|---|---|
| pPP3445 | CEN ARS TRP1 $P_{GAL1}$-mCherry-ste5(Δ1-214/VASP)-CTM | this study |
| pPP3446 | CEN ARS TRP1 $P_{GAL1}$-mCherry-ste5(370-590)-CTM | this study |
| pPP3447 | CEN ARS URA3 $P_{GAL1}$-mYFP-ste5(Δ1-214/I504T)-CTM | this study |
| pPP3448 | CEN ARS URA3 $P_{GAL1}$-mCherry-ste5(Δ1-214/I504T)-CTM | this study |
| pPP3449 | CEN ARS URA3 $P_{GAL1}$-mYFP-ste5(560-820)-CTM | this study |
| pPP3450 | CEN ARS URA3 $P_{GAL1}$-mCherry-ste5(560-820)-CTM | this study |
| pPP3451 | CEN ARS TRP1 $P_{GAL1}$-mYFP-ste5(370-590)-CTM | this study |
| pPP3469 | CEN ARS TRP1 $P_{STE5}$-ste5(1-560/ΔPM)-$HA_3$ $T_{CYC1}$ | this study |
| pPP3470 | CEN ARS TRP1 $P_{STE5}$-ste5(1-560/ΔRING)-$HA_3$ $T_{CYC1}$ | this study |
| pPP3471 | CEN ARS URA3 $P_{STE5}$-ste5(I504T/ΔPM)-$myc_{13}$ $T_{CYC1}$ | this study |
| pPP3472 | CEN ARS URA3 $P_{STE5}$-ste5(I504T/ΔRING)-$myc_{13}$ $T_{CYC1}$ | this study |
| pPP3473 | CEN ARS URA3 $P_{STE5}$-ste5(ΔPH::$PLC\delta^{PH}$/ΔPM)-$myc_{13}$ $T_{CYC1}$ | this study |
| pPP3474 | CEN ARS URA3 $P_{STE5}$-ste5(ΔPH::$PLC\delta^{PH}$/ΔRING)-$myc_{13}$ $T_{CYC1}$ | this study |
| pPP3514 | CEN ARS TRP1 $P_{CYC1}$-ste5[N]-ste11[C] $T_{ADH1}$ | this study |
| pPP3529 | CEN ARS TRP1 $P_{CYC1}$-ste20[N]-ste11[C] $T_{ADH1}$ | this study |
| pPP3565 | CEN ARS URA3 $P_{STE5}$-ste5(I504T/PH*)-$myc_{13}$ $T_{CYC1}$ | this study |
| pPP3578 | CEN ARS TRP1 $P_{CYC1}$-ste20[N]-ste11[C] $T_{ADH1}$ | this study |
| pPP3601 | CEN ARS TRP1 $P_{CYC1}$-(ste20[N]/CRIB*)-ste11[C] $T_{ADH1}$ | this study |
| pPP3602 | CEN ARS TRP1 $P_{CYC1}$-(ste20[N]/BR*)-ste11[C] $T_{ADH1}$ | this study |
| pPP3613 | CEN ARS URA3 $P_{CYC1}$-ste20[N]-ste5[VWA] $T_{ADH1}$ | this study |
| pPP3614 | CEN ARS URA3 $P_{CYC1}$-(ste20[N]/CRIB*)-ste5[VWA] $T_{ADH1}$ | this study |
| pPP3615 | CEN ARS URA3 $P_{CYC1}$-(ste20[N]/BR*)-ste5[VWA] $T_{ADH1}$ | this study |
| pPP3620 | CEN ARS TRP1 $P_{GAL1}$-GFP-ste5(V763A S861P)-CTM | this study |
| pPP3658 | CEN ARS URA3 $P_{GAL1}$-GFP-ste5(560-820/coa1)-CTM | this study |
| pPP3659 | CEN ARS URA3 $P_{GAL1}$-GFP-ste5(560-820/Δ7BR)-CTM | this study |
| pPP3660 | CEN ARS URA3 $P_{GAL1}$-GFP-ste5(560-820/coa2)-CTM | this study |
| pPP3661 | CEN ARS URA3 $P_{GAL1}$-GFP-ste5(560-820/coa3)-CTM | this study |
| pPP3670 | CEN ARS TRP1 $P_{STE5}$-ste5(1-531)-$HA_3$ $T_{CYC1}$ | this study |
| pPP3671 | CEN ARS TRP1 $P_{STE5}$-ste5(1-518)-$HA_3$ $T_{CYC1}$ | this study |
| pPP4408 | CEN ARS TRP1 $P_{CYC1}$-(ste5[N]/ΔPM)-ste11[C] $T_{ADH1}$ | this study |
| pPP4409 | CEN ARS TRP1 $P_{CYC1}$-(ste5[N]/ΔRING)-ste11[C] $T_{ADH1}$ | this study |
| pPP4410 | CEN ARS TRP1 $P_{CYC1}$-(ste5[N]/Gβγ*)-ste11[C] $T_{ADH1}$ | this study |
| pPP4411 | CEN ARS URA3 $P_{GAL1}$-GFP-ste5(590-820) | this study |
| pPP4412 | CEN ARS URA3 $P_{GAL1}$-GFP-ste5(Δ1-589) | this study |
| pRS314 | CEN ARS TRP1 vector | [S8] |
| pRS316 | CEN ARS URA3 vector | [S8] |
| pVMG110 | CEN ARS TRP1 STE7-$myc_{13}$ | [S9] |
| pVMG113 | CEN ARS TRP1 ste7(ND)-$myc_{13}$ | [S9] |

* mutations and sequence regions denoted by abbreviated names:

| | |
|---|---|
| Ste5 VASP | V763A S861P |
| Ste5 ΔPM | Δ48-67 |
| Ste5 ΔRING | Δ177-229 |
| Ste5 Gβγ* | Δ152-173 |
| Ste5 PH* | R407S K411S |
| Ste5 ΔPH | Δ364-532 |
| Ste5[N] | aa 2-281 |
| Ste5[VWA] | aa 560-820 |
| Ste5 coa1 | N744A D746A |
| Ste5 coa2 | V745A Y747A S748A |
| Ste5 coa3 | K750A D752A E753A |
| Ste5 Δ7BR | Δ778-788 |
| Ste20[N] | aa 2-403 |
| Ste20 CRIB* | S338A H345G |
| Ste20 BR* | K285N K286N R287G K288A R297A M298G K299A K310N R311G |
| Ste11-Asp3 | S281D S285D T286D |
| Ste11[C] | aa 117-717 |
| Ste7 ND | R9A R10A L17A L19A R62A R63A L69A L71A |
| Ste7 EE | S359E T363E |
| $PLC\delta^{PH}$ | aa 11-140 |

### SUPPLEMENTAL FIGURE LEGENDS

**FIGURE S1:** *Trans* signaling efficiency and Ste5 sequence requirements (related to Figure 2).

(A) *Top*, transcriptional induction (*FUS1-lacZ*) in strains (PPY1974, PPY1975, PPY2032) with the indicated *STE5* allele harboring Ste5-myc mutants on plasmids (as in Figure 1C), after treatment with α factor (5 μM, 2 hr). Bars, mean ± SD (n = 3). *Bottom*, example of levels of Ste5-myc protein in the same left-to-right order as in the bar graph above; G6PDH served as a loading control.

(B) *Top*, patch mating assay of *ste5Δ* and *ste5-I504T* strains (PPY2032, PPY1975) harboring *STE5-HA₃* plasmids (as in Figure 1D,E). *Bottom*, example showing protein levels for the Ste5-HA variants used above (in PPY1975), from experiments as shown in Figure 1E.

(C) Comparison of pathway output (*FUS1-lacZ*) in cells with wild-type Ste5 (WT) compared to cells expressing complementing Ste5 mutants, tested in two different strain backgrounds (S288C and W303). Strains: PPY1955, PPY1974, PPY1975, PPY640, PPY1967, PPY1968. Bars: mean ± range (n = 2). The complemented levels were roughly 40% of WT in the S288C strains, or 20-30% in the W303 strains.

**FIGURE S2:** Domain requirements for *trans* signaling (related to Figure 3).

(A) Comparison of functionally defined domains in Ste5 with predicted globular and disordered regions. *Top*, plot of disorder probability consensus (i.e., the mean of five prediction algorithms: PONDR-VLXT, PONDR-VSL2, IUpred-L, ANCHOR, and ESpritz-N; see [S10]). *Middle*, plot of predicted globular regions using GlobPlot [S11].

(B) Truncations define the C-terminal end of PH domain required for *trans* signaling in response to pheromone rather than membrane-tethering. Strains (PPY1975, PPY2032) harboring *STE5-HA₃* plasmids were assayed for *FUS1-lacZ* induction (mean ± SD; n = 4) in response to α factor (5 μM, 2 hr). The diagram at top shows the truncation endpoints.

(C) *Trans* signaling by Ste5-CTM variants with different fluorescent protein tags. Ste5ΔN-CTM constructs (left) or minimal PH-CTM and VWA-CTM constructs (right) included three different N-terminal fluorophore tags: enhanced GFP (S65A V68L S72A), monomeric YFP (mYFP: S65G V68L S72A Q80R T203Y A206K K238N), and monomeric Cherry (mCherry; Genbank sequence AY678264). All coexpression permutations were capable of inducing *FUS1-lacZ* (mean ± range; n = 2). Strain: PPY886.

**FIGURE S3:** Ste5 function tolerates swapping of PH and VWA domains (related to Figure 4).

*Top*, in Ste5 “RVP” (RING-VWA-PH) the positions of the PH and VWA domains are interchanged. *Bottom*, signaling assays of Ste5 variants lacking individual domains or containing swapped domains (on *STE5-myc$_{13}$* plasmids). *FUS1-lacZ* induction (mean ± SD) was measured after treatment of cells (PPY2032) with $\alpha$ factor (5 µM, 2 hr; n = 3) or after galactose treatment (3 hr; n = 6) in cells (PPY886) with *P$_{GAL1}$-STE4* or *P$_{GAL1}$-STE11-Cpr*.

**FIGURE S4:** Analysis of Ste5-I504T variants, Ste7 mutants, and Ste5 VWA domain mutants (related to Figure 5).

(A) Experiment as in Figure 5B, but here the *P$_{GAL1}$-STE5-CTM* and *P$_{GAL1}$-STE11-Cpr* constructs were induced at full strength using galactose (2 hr); under these conditions the small VWA fragment (590-820) shows reduced dependence on the CTM tether. Strain: PPY886 (mean ± SD, n = 3).

(B) MAPK phosphorylation in strains coexpressing *STE5-myc$_{13}$* variants with different complementing partners. Cells (PPY2032) with Ste5-VASP were treated ± $\alpha$ factor (5 µM, 15 min). Cells (PPY858) with *P$_{GAL1}$*-expressed Ste11-Cpr or Ste5 PH-CTM were treated with galactose for 90 min. and then incubated ± $\alpha$ factor (5 µM, 30 min.). At top, a representative example of Ste5-myc levels is shown.

(C) Ste7 mutants do not complement each other. *Top*, a *ste7*$\Delta$ strain (PPY891) harbored plasmid-borne Ste7 mutants, including Ste7-ND (a “non-docking” mutant, with mutations in two Fus3-binding motifs [S9]) and Ste7-R220 (a catalytically inactive mutant; [S12]). *FUS1-lacZ* induction (mean ± SD, n = 3) was measured ± $\alpha$ factor (5 µM, 2 hr). *Bottom*, schematic depiction of the result.

(D) Ste5 VWA domain mutations affecting the Ste7 → Fus3 reaction. Regions (i) and (ii) denote the coactivator loop (responsible for “catalytically unlocking” Fus3) and the Ste7-binding region [S13], respectively, as highlighted in the structure (PDB ID: 3FZE). Sequence changes are shown for three coactivator loop mutants (coa1-3) and a deletion of the Ste7-binding region ($\Delta$7BR). Mutants coa1 and $\Delta$7BR are identical to “mut C” and “mut B”, respectively, which were extensively probed for effects on the Ste7 → Fus3 reaction in vitro [S13]; (coa2 and coa3 are related to mutants 14 and 15 from that study).

(E) *Top*, *FUS1-lacZ* induction (mean ± SD, n = 3) for *STE5-myc$_{13}$* variants, tested in both *kss1*$\Delta$ (PPY1669) and *KSS1* (PPY858) contexts (5 µM $\alpha$ factor, 2 hr) to ensure that the results reflect Fus3 activation. Note the mild defects of the VWA mutants, compared to Ste5-VASP.

*Bottom*, the same mutants were tested for MAPK phosphorylation (5 µM $\alpha$ factor, 15 min.); a non-specific band on the same blot served as a loading control.

(F) Mutant forms of a VWA-CTM construct were coexpressed with PH-CTM, Ste20[N]-Ste11[C], and Ste7-EE, a pseudo-active form of Ste7 that was the source of Ste7 kinase activity for prior in vitro assays [S13]. *FUS1-lacZ* induction (mean ± SD, n = 3) was measured in *ste5*Δ *kss1*Δ cells (PPY1669) after galactose treatment (3 hr).

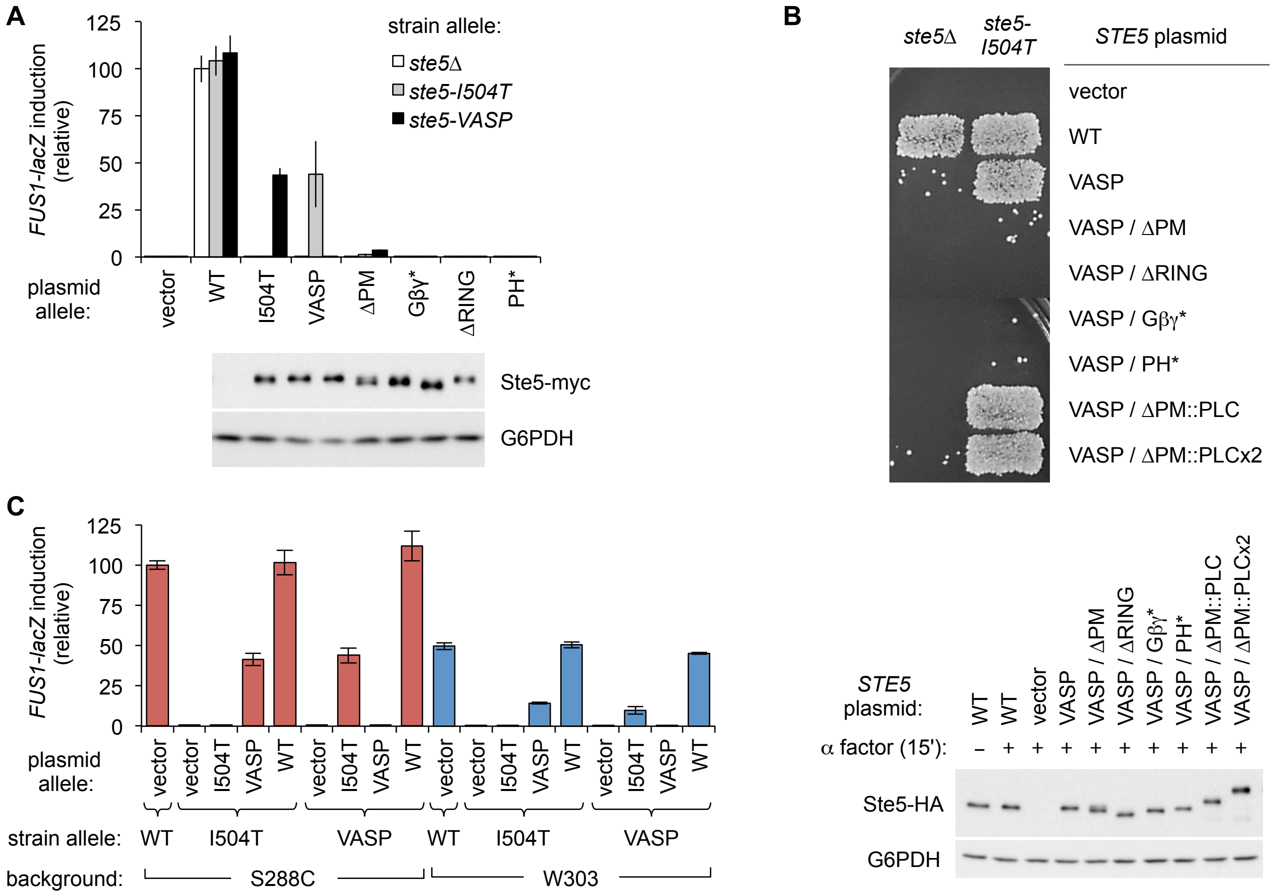


**FIGURE S1:** *Trans* signaling efficiency and Ste5 sequence requirements (related to Figure 2).

(A) *Top*, transcriptional induction (*FUS1-lacZ*) in strains (PPY1974, PPY1975, PPY2032) with the indicated *STE5* allele harboring Ste5-myc mutants on plasmids (as in Figure 1C), after treatment with α factor (5 μM, 2 hr). Bars, mean ± SD (n = 3). *Bottom*, example of levels of Ste5-myc protein in the same left-to-right order as in the bar graph above; G6PDH served as a loading control.

(B) *Top*, patch mating assay of *ste5*Δ and *ste5-I504T* strains (PPY2032, PPY1975) harboring *STE5-HA$_3$* plasmids (as in Figure 1D,E). *Bottom*, example showing protein levels for the Ste5-HA variants used above (in PPY1975), from experiments as shown in Figure 1E.

(C) Comparison of pathway output (*FUS1-lacZ*) in cells with wild-type Ste5 (WT) compared to cells expressing complementing Ste5 mutants, tested in two different strain backgrounds (S288C and W303). Strains: PPY1955, PPY1974, PPY1975, PPY640, PPY1967, PPY1968. Bars: mean ± range (n = 2). The complemented levels were roughly 40% of WT in the S288C strains, or 20-30% in the W303 strains.

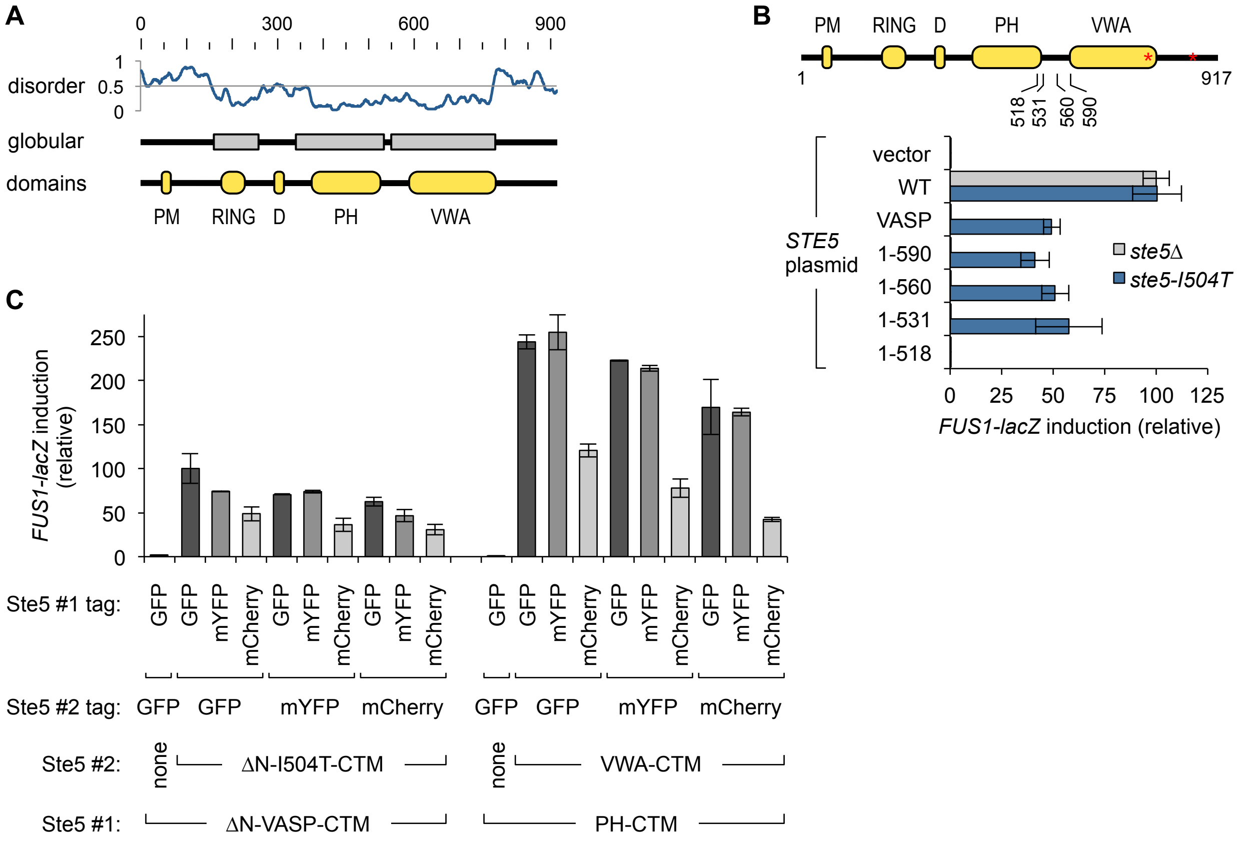


**FIGURE S2:** Domain requirements for *trans* signaling (related to Figure 3).

(A) Comparison of functionally defined domains in Ste5 with predicted globular and disordered regions. *Top*, plot of disorder probability consensus (i.e., the mean of five prediction algorithms: PONDR-VLXT, PONDR-VSL2, IUpred-L, ANCHOR, and ESpritz-N; see [S10]). *Middle*, plot of predicted globular regions using GlobPlot [S11].

(B) Truncations define the C-terminal end of PH domain required for *trans* signaling in response to pheromone rather than membrane-tethering. Strains (PPY1975, PPY2032) harboring *STE5-HA*$_3$ plasmids were assayed for *FUS1-lacZ* induction (mean ± SD; n = 4) in response to $\alpha$ factor (5 µM, 2 hr). The diagram at top shows the truncation endpoints.

(C) *Trans* signaling by Ste5-CTM variants with different fluorescent protein tags. Ste5ΔN-CTM constructs (left) or minimal PH-CTM and VWA-CTM constructs (right) included three different N-terminal fluorophore tags: enhanced GFP (S65A V68L S72A), monomeric YFP (mYFP: S65G V68L S72A Q80R T203Y A206K K238N), and monomeric Cherry (mCherry; Genbank sequence AY678264). All coexpression permutations were capable of inducing *FUS1-lacZ* (mean ± range; n = 2). Strain: PPY886.

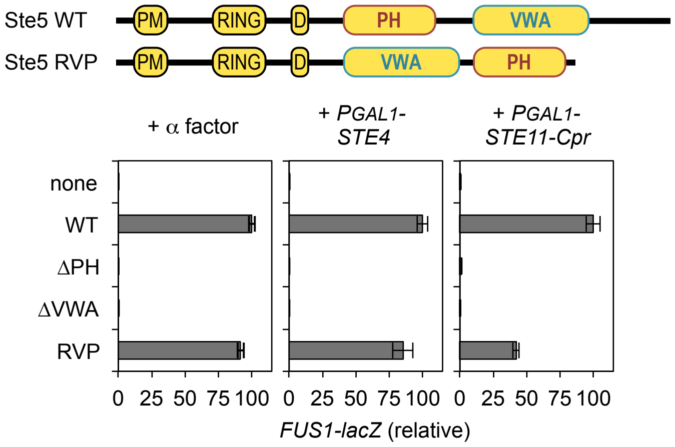


**FIGURE S3:** Ste5 function tolerates swapping of PH and VWA domains (related to Figure 4).
*Top*, in Ste5 "RVP" (RING-VWA-PH) the positions of the PH and VWA domains are interchanged. *Bottom*, signaling assays of Ste5 variants lacking individual domains or containing swapped domains (on *STE5-myc*$_{13}$ plasmids). *FUS1-lacZ* induction (mean ± SD) was measured after treatment of cells (PPY2032) with α factor (5 μM, 2 hr; n = 3) or after galactose treatment (3 hr; n = 6) in cells (PPY886) with $P_{GAL1}$-*STE4* or $P_{GAL1}$-*STE11-Cpr*.

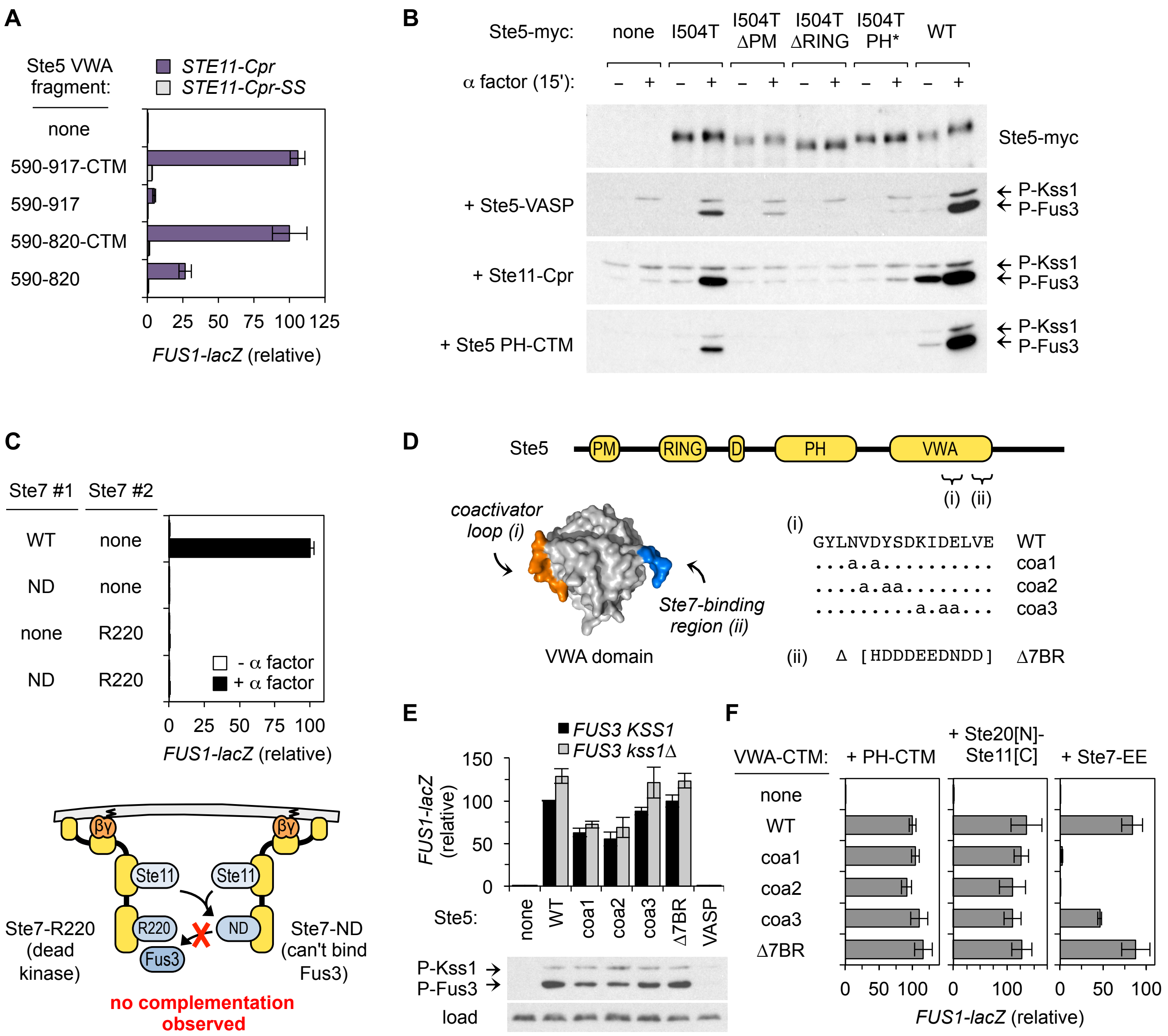


**FIGURE S4:** Analysis of Ste5-I504T variants, Ste7 mutants, and Ste5 VWA domain mutants (related to Figure 5).

(A) Experiment as in Figure 5B, but here the *$P_{GAL1}$-STE5-CTM* and *$P_{GAL1}$-STE11-Cpr* constructs were induced at full strength using galactose (2 hr); under these conditions the small VWA fragment (590-820) shows reduced dependence on the CTM tether. Strain: PPY886 (mean ± SD, n = 3).

(B) MAPK phosphorylation in strains coexpressing *$STE5$-$myc_{13}$* variants with different complementing partners. Cells (PPY2032) with Ste5-VASP were treated ± α factor (5 μM, 15 min). Cells (PPY858) with $P_{GAL1}$-expressed Ste11-Cpr or Ste5 PH-CTM were treated with galactose for 90 min. and then incubated ± α factor (5 μM, 30 min.). At top, a representative example of Ste5-myc levels is shown.

(C) Ste7 mutants do not complement each other. *Top*, a *ste7Δ* strain (PPY891) harbored plasmid-borne Ste7 mutants, including Ste7-ND (a "non-docking" mutant, with mutations in two Fus3-binding motifs [S9]) and Ste7-R220 (a catalytically inactive mutant; [S12]). *FUS1-lacZ* induction (mean ± SD, n = 3) was measured ± α factor (5 μM, 2 hr). *Bottom*, schematic depiction of the result.

(D) Ste5 VWA domain mutations affecting the Ste7 → Fus3 reaction. Regions (i) and (ii) denote the coactivator loop (responsible for "catalytically unlocking" Fus3) and the Ste7-binding region [S13], respectively, as highlighted in the structure (PDB ID: 3FZE). Sequence changes are shown for three coactivator loop mutants (coa1-3) and a deletion of the Ste7-binding region (Δ7BR). Mutants coa1 and Δ7BR are identical to "mut C" and "mut B", respectively, which were extensively probed for effects on the Ste7 → Fus3 reaction in vitro [S13]; (coa2 and coa3 are related to mutants 14 and 15 from that study).

(E) *Top*, *FUS1-lacZ* induction (mean ± SD, n = 3) for *$STE5$-$myc_{13}$* variants, tested in both *kss1Δ* (PPY1669) and *KSS1* (PPY858) contexts (5 μM α factor, 2 hr) to ensure that the results reflect Fus3 activation. Note the mild defects of the VWA mutants, compared to Ste5-VASP. *Bottom*, the same mutants were tested for MAPK phosphorylation (5 μM α factor, 15 min.); a non-specific band on the same blot served as a loading control.

(F) Mutant forms of a VWA-CTM construct were coexpressed with PH-CTM, Ste20[N]-Ste11[C], and Ste7-EE, a pseudo-active form of Ste7 that was the source of Ste7 kinase activity for prior in vitro assays [S13]. *FUS1-lacZ* induction (mean ± SD, n = 3) was measured in *ste5Δ kss1Δ* cells (PPY1669) after galactose treatment (3 hr).
